# Introducing Isotòpia: a stable isotope database for Classical Antiquity

**DOI:** 10.1101/2023.10.19.563074

**Authors:** Giulia Formichella, Silvia Soncin, Carmine Lubritto, Mary Anne Tafuri, Ricardo Fernandes, Carlo Cocozza

## Abstract

We present Isotòpia, an open-access database compiling over 36,000 stable isotope measurements (δ^13^C, δ^15^N, δ^18^O, δ^34^S, ^87^Sr/^86^Sr, ^206^Pb/^204^Pb, ^207^Pb/^204^Pb, ^208^Pb/^204^Pb, ^207^Pb/^206^Pb, and ^208^Pb/^206^Pb) on human, animal, and plant bioarchaeological remains dating to Classical Antiquity (approximately 800 BCE - 500 CE). These were recovered from different European regions, particularly from the Mediterranean. Isotòpia provides a comprehensive characterisation of the isotopic data, encompassing various historical, archaeological, biological, and environmental variables. Isotòpia is a resource for meta-analytical research of past human activities and paleoenvironments. The database highlights data gaps in isotopic classical archaeology, such as the limited number of isotopic measurements available for plants and animals, limited number of studies on spatial mobility, and spatial heterogeneity of isotopic research. As such, we emphasise the necessity to address and fill these gaps in order to unlock the reuse potential of this database.

## Introduction

Classical Antiquity (c. 800 BCE - 500 CE) is a formative period for the Western world, and beyond. Greek and Roman civilisations, together with other Iron Age cultures left a lasting imprint on European material and immaterial culture [1–5]. The development of a ’Graeco-Roman’ culture was a complex process involving interactions among various peoples and civilizations [6–8]. Our knowledge of this culture has primarily hinged on the study of surviving Greek and Latin written documents or of artworks and monuments. These offer a narrow perspective, biased towards the lifeways of the ruling class and missing out on the broader population [9].

Isotopic methods offer direct information on the lifeways of past individuals through the analysis of their remains and make an important contribution to reduce research bias [10, 11]. Recently, the application of stable isotope measurements on bioarchaeological remains has significantly increased [12]. Isotopes are now employed in a variety of archaeological research questions, including the study of subsistence practices, spatial mobility, infant feeding strategies, and human-environment interactions [13–49]. This research potential has led to an exponentially increase of isotopic applications, and to recent efforts towards ’Big Isotopic Data’ initiatives [50–55]. This novel approach operates at varying degrees of spatiotemporal resolution and has the potential to offer multilayered insights into past human and environmental phenomena [52, 56–58]. In combination with other sources of past evidence, large isotopic datasets may offer a wide range of historical information [52, 59].

We present Isotòpia [60], an open-access database that compiles stable carbon (δ^13^C), nitrogen (δ^15^N), oxygen (δ^18^O), and sulphur (δ^34^S) isotope data, along with radiogenic strontium (^87^Sr/^86^Sr) and lead (^206^Pb/^204^Pb, ^207^Pb/^204^Pb, ^208^Pb/^204^Pb, ^207^Pb/^206^Pb, and ^208^Pb/^206^Pb) ratio measurements from human, animal, and plant remains dating to Classical Antiquity. This resource is designed to enable future data meta-analyses researching past phenomena across a broad spectrum of historical, archaeological, biological, and environmental variables. Here we provided a detailed description of the database and highlight isotopic data gaps in Classical Archaeology. Addressing these gaps could significantly enhance our knowledge resolution on the spatiotemporal dynamics of human and environmental systems throughout antiquity.

### Collection criteria for Isotòpia

Isotòpia collects isotopic measurements (δ^13^C, δ^15^N, δ^18^O, δ^34^S, ^87^Sr/^86^Sr, ^206^Pb/^204^Pb, ^207^Pb/^204^Pb, ^208^Pb/^204^Pb, ^207^Pb/^206^Pb and ^208^Pb/^206^Pb) and associated archaeological, historical, biological, and environmental information. Collected isotopic data was obtained from human or animal osteological materials or plant organic remains dated between 800 BCE and 500 CE; these were recovered from a territory matching the maximum geographical extent of the Roman Empire. However, in the case of Greece, the temporal range was extended to 3200 BCE to include isotopic data from Bronze Age civilizations [61–63].

The collection of isotopic measurements was carried out between November 2021 and March 2023 and relied on scholarly publications such as journal articles, archaeological reports, book chapters and academic dissertations written in English, Italian, Spanish, German and French. These texts were located online through the use of search engines and databases such as Google Scholar, Scopus, Researchgate.net and Academia.edu, and employing keywords such as ’isotopes’, ’human/faunal/plant remains’ and cultural and geographic tags (e.g., ’Roman’, ’Greece’). Missing publications were also identified by consulting the bibliography of the recovered texts. Metadata was retrieved from the same isotopic publications or, occasionally, from related non-isotopic works.

### Design and structure of Isotòpia

Isotòpia is a database designed to promote the reuse and interpretation of isotopic data for historical research. The metadata structure of the database includes multiple archaeological, historical, biological, and environmental variables. Furthermore, Isotòpia identifies sources of retrieved data. A comprehensive description of the metadata structure can be found in S1 Table.

Isotòpia is organised into three main tables listing human, animal, and plant data. There are some small differences in metadata structure among these. Each table contains a unique integer number sequence, and each number corresponds to one or more isotopic measurements for a registered sample. To allow for tracking and referencing of registered isotopic measurements with original sources, multiple identification codes (e.g. sample and individual IDs) are provided.

Reported conventions of biological sex and age-at-death may differ across publications. When necessary, we converted these according to a modified Buikstra and Ubelaker’s (1994) [64] classification (further information in S1 Appendix). This classification was based on the estimated age-at-death range retrieved from the original source. Whenever this range was encompassing more classes, these were reported and separated by a semicolon.

Isotòpia is a georeferenced database. In addition to contemporary administrative divisions, geographical coordinates based on the WGS84 reference system are documented to locate the locations of archaeological sites where samples were collected. In cases where coordinates were not published in the original data source, these were obtained using Google Earth, and an estimated radius of uncertainty for site location is given in the database. Additionally, the database includes a brief description of the archaeological site, its environmental features, and historical information.

Each entry is assigned a chronological range using the BCE/CE convention, with negative numbers representing years BCE. Also described are dating methods. When based on radiocarbon measurements, uncalibrated ^14^C dates (BP) and uncertainties plus radiocarbon lab codes are reported. Recognising that traditional periodisation systems for Classical Antiquity cannot be applied to all contexts included in Isotòpia, we developed a standardised scheme that merges together traditional Greek and Roman historical timelines (Table 1). Other periodisations, whenever reported in isotopic publications, are registered separately. Furthermore, the database includes descriptions of burial contexts, individuals’ social status, and indications of Christian religious affiliation.

**Table 1.** Periodisation used in Isotòpia.

| Broad Classification | Category used in Isotòpia | Date range BCE/CE | Target |
| --- | --- | --- | --- |
| Greek Bronze and Iron Age | Helladic | 3200 BCE – 1100 BCE | Bronze Age samples from Greece not found in Minoan and Mycenaean contexts |
|  | Minoan | 2700 BCE – 1400 BCE | Samples from the Isle of Crete |
|  | Mycenaean | 1750 BCE – 1100 BCE | Samples from Greece found in Mycenaean contexts, including those from the Isle of Crete dated 1450 BCE - 1100 BCE |
|  | Protogeometric | 1100 BCE – 900 BCE | Samples from Greece |
|  | Geometric | 900 BCE – 800 BCE | Samples from Greece |
| Greek | Archaic | 800 BCE – 480 BCE | All samples |
|  | Classical | 480 BCE – 323 BCE | All samples |
|  | Hellenistic | 323 BCE – 31 BCE | All samples |
| Roman Empire | Imperial | 31 BCE – 284 CE | All samples |
|  | Late Imperial | 284 CE – 476 CE | All samples |
| Post-Roman | Medieval | after 476 CE | All samples |

Isotòpia includes a full bibliographic reference reported according to APA guidelines, link to the publication website, and DOIs for the original data sources. In alignment with its collaborative design, Isotòpia also includes fields that cite other isotopic databases, whenever data was collected from previous compilations [52, 59, 65].

Stable isotope ratios employed in archaeological research are more commonly measured in organic (collagen protein) or biomineral (bioapatite/enamel) samples obtained from skeletal remains of humans and animals, or from charred plant remains. In Isotòpia, each entry can contain one or more stable isotope measurements and their associated uncertainties for a sample. Wherever original publications only provided mean population isotopic values, these are entered in the database as a single measurement together with the number of samples used to calculate the mean.

Isotòpia distinguishes carbon stable isotope measurements made using Isotope Ratio Mass Spectrometers (IRMS) and Accelerator Mass Spectrometers (AMS). This latter is used in the radiocarbon dating method and δ^13^C measurements from the two systems are often not comparable [66, 67]. Therefore, Isotòpia includes separate AMS and IRMS fields for δ^13^C and their associated uncertainties. Within Isotòpia, skeletal human and animal δ^13^C values measurements are reported for organic (collagen protein) and biomineral (bioapatite/enamel) components. These dietary proxies are determined by separate metabolic pathways and thus provide complementary dietary information [68].

For spatial mobility analysis, Isotòpia includes stable oxygen isotope measurements of carbonate or phosphate fractions of bioapatite [69, 70]. Moreover, δ^18^O_carbonate_ and δ^18^O_phosphate_ values can be reported relative to the Vienna Pee-Dee-Belemnite (VPDB) or Vienna Standard Mean Ocean Water (VSMOW) analytical standards. All these combinations are considered in Isotòpia. These values can be converted into estimates of δ^18^O ingested water [71–73]. Given that some publications only reported calculated δ^18^O_drinking-water_ values, Isotòpia includes a field for these.

Different collagen preservation criteria have been proposed to assess the reliability of isotopic measurements [74–77]. Collagen yield, %C, %N, %S, atomic C:N ratio, atomic C:S ratio, and atomic N:S ratio values are included in Isotòpia when provided in the original publication. As acceptable ranges for these criteria may differ across different publications, the database allows users to select which criteria to employ.

### Isotòpia access and reuse potential

Isotòpia human, animal, and plant datasets can be accessed in both EXCEL and CSV formats at https://www.doi.org/10.48493/m0m0-b436 [60]. The database is stored within *AENEAS: a bioarchAEological data commuNity for GraEco-RomAn claSsical antiquity* (https://pandoradata.earth/organization/aeneas-a-bioarchaeological-data-community-for-greco-roman-classical-antiquity). The AENEAS data community is hosted by the Pandora data platform (https://pandoradata.earth/). AENEAS was designed to compile datasets pertaining to Classical Antiquity. Isotòpia is also part of the IsoMemo network (https://isomemo.com/), an open-access collaboration of independent isotopic databases. Isotòpia shares the collaborative and distributive ethos of the network. In this context, Isotòpia also adheres to the FAIR and CARE principles, an established set of guidelines for effective scientific data management [78, 79].

A total of 11,205 human, 4,707 animal, and 934 plant entries were amassed from 281 primary sources. S2 Appendix provides a comprehensive list of publications from where isotopic data was retrieved. The database contains a total of 36,922 isotopic measurements for δ^13^C, δ^15^N, δ^18^O, δ^34^S, ^87^Sr/^86^Sr, ^206^Pb/^204^Pb, ^207^Pb/^204^Pb, ^208^Pb/^204^Pb, ^207^Pb/^206^Pb, and ^208^Pb/^206^Pb. The spatial distribution of the entries is shown in Figure 1 and Figure 2 illustrates the temporal distribution of the investigated sites. Data records are illustrated in S3 Appendix.

**Fig 1.**
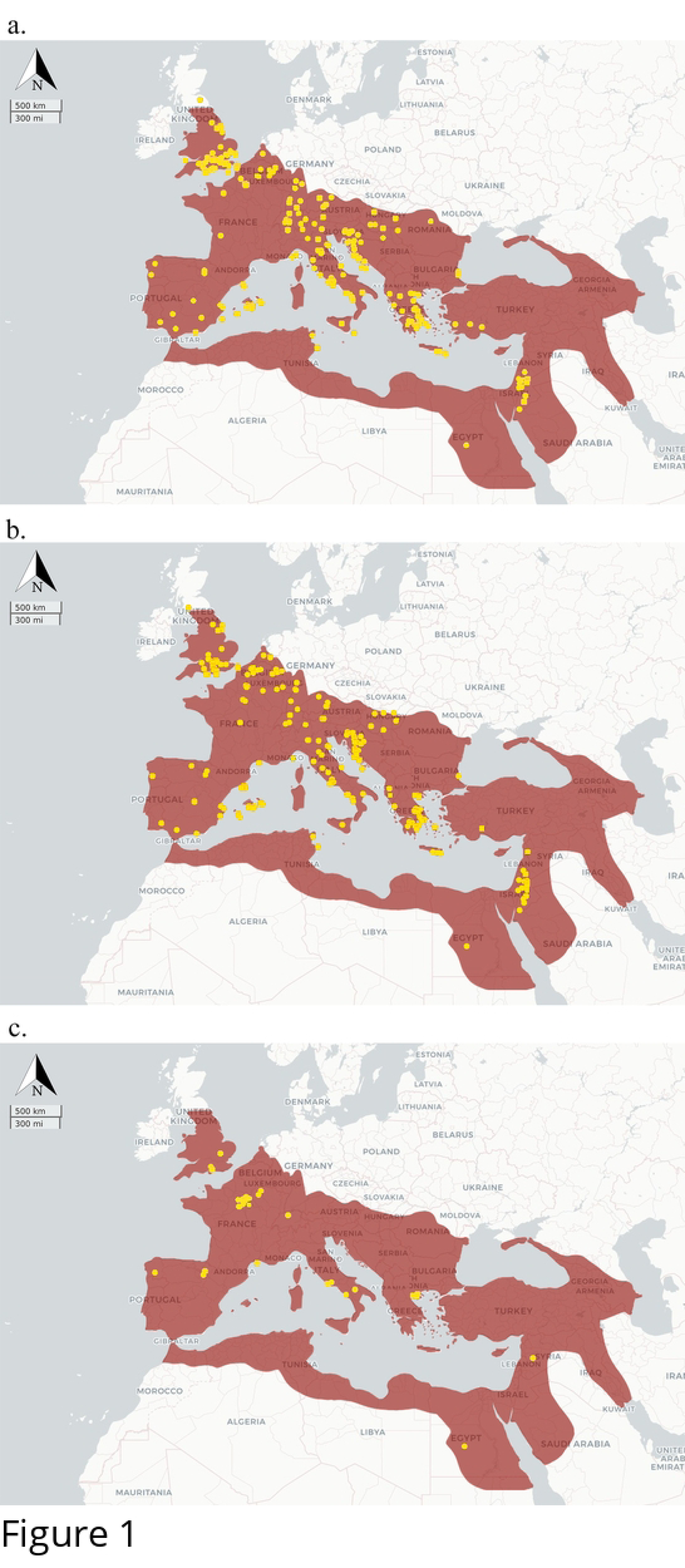
Spatial distribution of sites registered in Isotòpia for a) humans; b) animals; c) plants.

**Fig 2.**
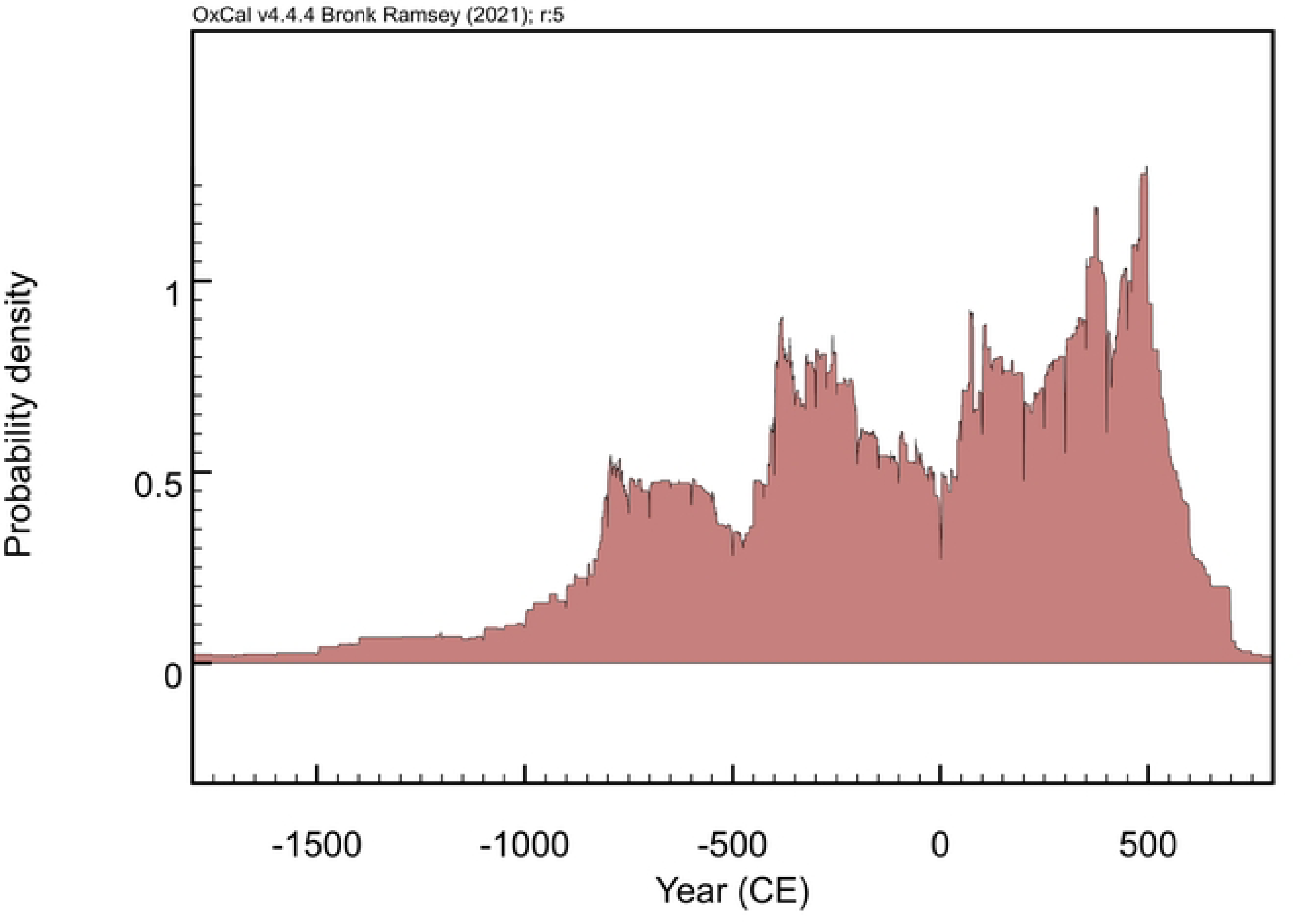
Temporal density of data collected in Isotòpia. Plot produced using OxCal 4.4.4. [80].

Isotòpia detailed metadata structure and isotopic data offer many possibilities for reuse. This can be expanded by combining Isotòpia with other databases via the Pandora platform. Research applications include reconstructing past climates and environments and the interplay between these and humans, tracking the development of farming practices, and investigating past human dietary and spatial mobility patterns. Moreover, as archaeological sites are spatiotemporally referenced, Isotòpia can be employed as a catalogue for heritage management. Figure 3 presents a schematic diagram delineating Isotòpia’s repository architecture and potential research applications.

**Fig 3.**
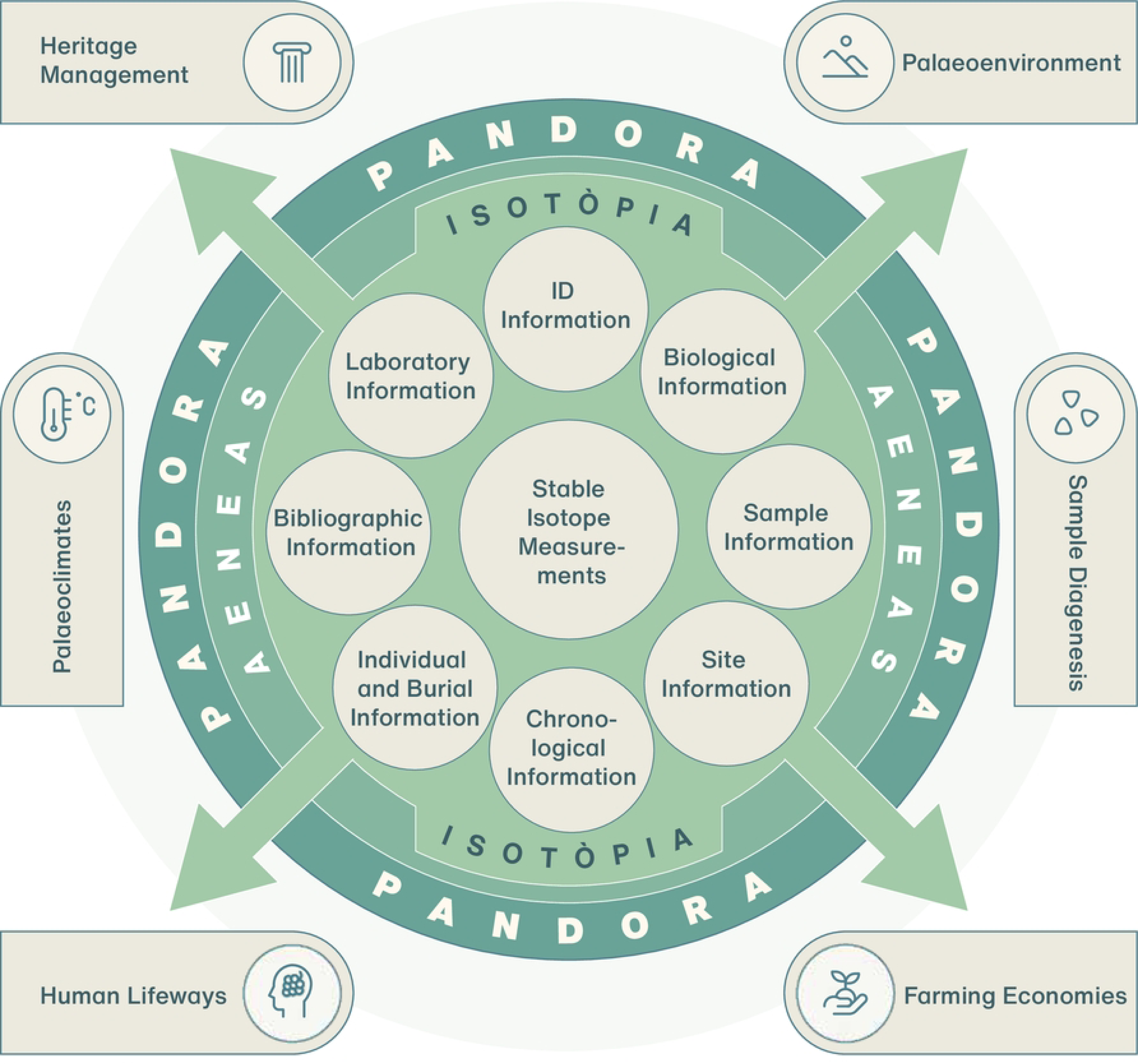
Flowchart of Isotòpia repository architecture and potential research and heritage management applications. Design by Hans-Georg Sell for MPI GEA, 2023

### Isotopic research gaps in classical archaeology

Isotòpia can be used to identify data gaps in isotopic research and motivate future research. Such gaps can have several reasons. These include, greater research interest and funding availability to carry out projects in certain regions or time periods, spatial variations in preservation conditions, type of burial practices, among others.

Isotopic analyses of zooarchaeological and archaeobotanical remains are employed to establish isotopic baselines necessary to interpret human isotopic values used for the study of dietary and spatial mobility patterns. These materials are also used to reconstruct past agricultural practices and local environmental conditions [10, 81–83]. However, faunal and plant isotopic measurements are present in relatively small numbers. Human isotopic data represents 68.90% of the database, while zooarchaeological remains account for 26.43% and archaeobotanical samples for 4.67%. Beyond insufficient research on past farming economies and paleoenvironments, the low amount of animal and plant measurements also suggests that a substantial portion of isotopic research addressing human lifeways in Classical Antiquity lacks reference isotopic baselines.

The majority of isotopic values in Isotòpia consists of δ^13^C_collagen_ and δ^15^N_collagen_ measurements denoting a higher research interest in the study of past diets but also less diagenetic issues when compared to other human tissues, and lower measurement costs and labour intensity (Table 2).

**Table 2.** Number of isotopic measurements collected in Isotòpia according to each isotopic proxy.

|  | **Humans** | **Animals** | **Plants** | **Total** |
| --- | --- | --- | --- | --- |
| $\delta^{13}\text{C}_{\text{collagen}}$ | 8333 | 3980 | 1938 | 14251 |
| $\delta^{15}\text{N}_{\text{collagen}}$ | 7766 | 3976 | 1350 | 13092 |
| $\delta^{34}\text{S}_{\text{collagen}}$ | 598 | 84 | 0 | 682 |
| $\delta^{13}\text{C}_{\text{carbonate}}$ | 1556 | 641 | 0 | 2197 |
| $\delta^{18}\text{O}_{\text{carbonate}}$ (VPDB) | 1611 | 487 | 0 | 2098 |
| $\delta^{18}\text{O}_{\text{carbonate}}$ (VSMOW) | 515 | 24 | 0 | 539 |
| $\delta^{18}\text{O}_{\text{phosphate}}$ (VPDB) | 0 | 0 | 0 | 0 |
| $\delta^{18}\text{O}_{\text{phosphate}}$ (VSMOW) | 587 | 12 | 0 | 599 |
| $\delta^{18}\text{O}_{\text{drinking-water}}$<br>(VSMOW) | 82 | 8 | 0 | 90 |
| $^{87}\text{Sr}/^{86}\text{Sr}$ | 2204 | 780 | 0 | 2984 |
| $^{206}\text{Pb}/^{204}\text{Pb}$ | 78 | 0 | 0 | 78 |
| $^{207}\text{Pb}/^{204}\text{Pb}$ | 78 | 0 | 0 | 78 |
| $^{208}\text{Pb}/^{204}\text{Pb}$ | 78 | 0 | 0 | 78 |
| $^{207}\text{Pb}/^{206}\text{Pb}$ | 78 | 0 | 0 | 78 |
| $^{208}\text{Pb}/^{206}\text{Pb}$ | 78 | 0 | 0 | 78 |
| **Total** | 23642 | 9992 | 3288 | 36922 |

The Pandora&IsoMemo initiative offers open-access and user-friendly spatiotemporal R-based modelling tools for georeferenced datasets (https://isomemoapp.com/app/iso-memo-app). Within this we used the KernelTimeR options to group sites according to their spatial distribution and chronology. From this we evaluated temporal coverage for each group (Fig. 4). Modelling details are given in S4 Appendix.

**Fig 4.**
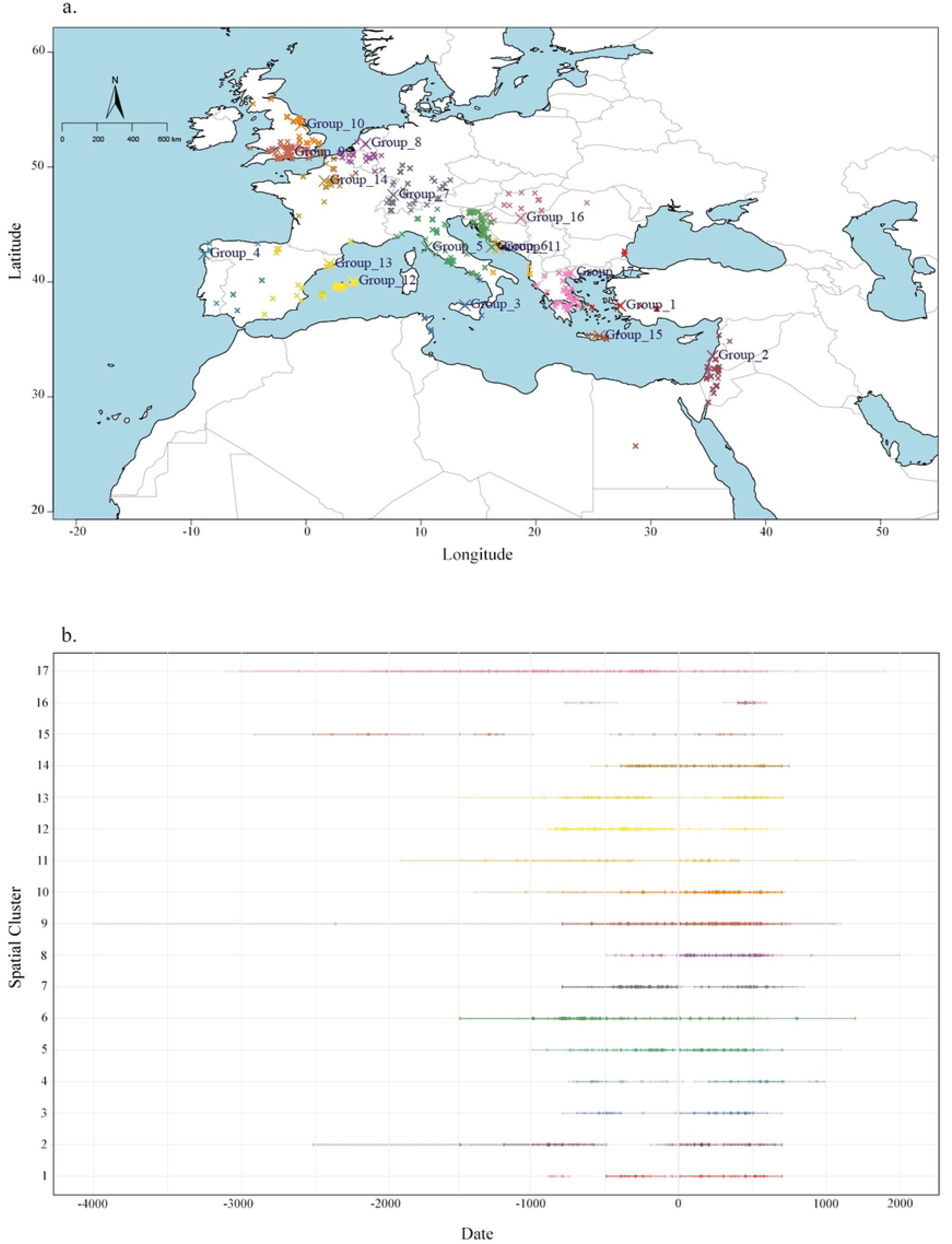
a: Modelled spatiotemporal groups. b: time ranges for each chronological range within the site according to group assignment.

Figure 4 shows that although North Africa and the eastern Mediterranean were zones of high economic and cultural significance they are poor in isotopic research [84, 85] (Fig. 4). Several factors might account for this, including a lower number of local/national isotopic laboratories and local environmental conditions which are more prone to sample degradation [74–77]. Western Mediterranean regions that served as critical nodes in cultural and commercial exchanges [86, 87], such as the islands of Corsica, Sardinia, and Sicily are also underrepresented. Conversely, the Balearic Islands show a higher density of isotopic data with the temporal distribution indicating that the majority of this data stems from studies on Phoenician and Roman cemeteries. The Iberian Peninsula has been subject to spatially dispersed isotopic research, with Portugal being largely underrepresented. Also France, despite its historical significance as ancient Gaul, exhibits a scarcity of isotopic data, save for the region surrounding Paris, where most studies are concentrated on pre-

Roman populations. Croatia and Slovenia display dense clusters of data, whereas the remainder of the Balkans and Central Europe are lacking isotopic data. Here, Hungary stands out as an exception. This broad spatiotemporal research gap, however, might be attributable to research traditions that place greater emphasis on other periods and historical events, such as group migrations during prehistoric and medieval periods [88–90].

The extensive compilation of data from Britain is a consequence of its deep-rooted tradition in osteological and biomolecular archaeological research. The temporal distribution of data here indicates a consistent employment of isotopic proxies, encompassing the Iron Age and Roman periods. Isotopic analyses have contributed to the study of Romanisation processes in Britain [13, 91]. The concentration of isotopic data in Greece and Italy is instead likely due to their historical significance as the cradles of ancient Greek and Roman civilizations. There is a particularly dense concentration of isotopic studies from Greece during the Classical (500-323 BCE) and Hellenistic (323-31 BCE) periods. Significant research focus also exists for Bronze and early Iron Age Greece, despite the challenges posed by larger temporal uncertainties. Conversely, there appears to be a comparatively smaller research focus on the Roman rule period in this region. Research on ancient Italy includes all periods, yet reveals a marked focus on the expansion era of the Roman Republic (third-second century BCE) and the Roman Empire. Despite the prevalent practice of cremation in the first century CE [92–94], which may render isotopic analysis impractical [95], the temporal distribution for Italy does not show a significant data gap. This might be mostly due to isotopic measurements generated from Pompeii and Herculaneum, two archaeological sites preserved by the 79 CE eruption of Mount Vesuvius [33, 34, 96, 97]. However, there is also additional data from other sites that presents larger temporal uncertainty. To improve the resolution of future investigations, we therefore advocate further direct dating methodologies, such as radiocarbon analysis, to be modelled together with the available historical and archaeological evidence.

## Conclusion

The Isotòpia open access database compiles c. 36,000 isotopic measurements on human, animal, and plant bioarchaeological samples dating to Classical Antiquity. This extensive collection allows for various research applications at different spatiotemporal scales of study within the disciplines of history, archaeology, physical anthropology, stable isotopes, and palaeoecology. In an effort to shape forthcoming research trajectories, we highlighted the presence of isotopic data gaps across materials, proxies, regions, and time periods. This revealed a scarcity of isotopic measurements for faunal and plant remains, an inclination towards dietary studies utilising stable carbon and nitrogen isotope analysis, and a preponderance of studies focusing on Classical and Hellenistic Greece, Roman Italy, and the Romanisation process in Britain. We thus propose that future research initiatives should be undertaken outside these contexts, as this would improve the current knowledge on the lifeways of ancient populations during Classical Antiquity and of the environments that surrounded them.

## Acknowledgments

The data was collected as part of the Pandora & IsoMemo initiatives supported by Max Planck Institute of geoanthropology, PS&H research group, University of Warsaw, Masaryk University, and Eurasia3angle research group. We thank Martina Farese for sharing the data collected in the MAIA database before its publication. We Thank Hans-Georg Sell for Hans-Georg Sell for creating the flowchart of Isotòpia. We also thank all researchers who have published stable isotope measurements that fit with the scopes of this database and contributed indirectly to the formation of this collection.

## Supporting information captions

S1 Table: Description of Isotòpia’s metadata structure

S2 Appendix: List of scientific publications from which isotopic data is retrieved

S3 Appendix: Summary statistics and data records for Isotòpia

S4 Appendix: Spatiotemporal groupings using KernelTimeR

## CRediT Statement

Giulia Formichella: Design of metadata structure (equal); Data Collection (lead); Writing - original draft (lead); Visualisation (lead); Formal analysis (lead).

Silvia Soncin: Conceptualisation (equal); Data Collection (supporting); Supervision (equal); Writing – original draft (supporting); Writing – review and editing (lead).

Carmine Lubritto: Writing – review and editing (equal).

Mary Anne Tafuri: Supervision (equal); Writing – review and editing (equal).

Ricardo Fernandes: Conceptualisation (lead); Methodology (lead); Design of metadata structure (equal); Writing – review and editing (supporting); Funding Acquisition (lead).

Carlo Cocozza: Conceptualisation (lead); Design of metadata structure (equal); Data Collection (supporting); Supervision (lead); Writing – original draft (supporting); Writing – review and editing (lead); Visualisation (supporting); Formal analysis (supporting).

